# Comparative antioxidant activity of *Ocimum gratissimum* and *Jatropha tanjorensis* 2024

**DOI:** 10.1101/2024.10.30.620967

**Authors:** Emmanuel Chukwuebuka Ike, Paschal N. Wokota, Lynda Nnenna Gabez

## Abstract

**Introduction:** The leaves of *Ocimum gratissimum* and *Jatropha tanjorensis* are widely used together as vegetables in food and in several preparations for medicinal purposes. For this reason, their combined antioxidant potentials were evaluated to ascertain if such preparations should be encouraged or otherwise in relation to their individual antioxidant effects.

**Materials and Methods:** the combined extract of *O.gratissimum* and *J. tanjorensis* (CEOJ) were serially diluted into different concentrations; concentrations of 1.9µg/ml, 3.91µg/ml, 7.81µg/ml, 15.63µg/ml, 31.25µg/ml, 62.5µg/ml, 125µg/ml and 250µg/ml were made and used for the assays. the single extracts were also diluted into similar concentrations. DPPH in methanol was used as blank for the DPPH-free radical scavenging assay. The absorbance of the concentrations and that of the controls were read at 517nm and 562nm for the DPPH assay using a spectrophotometer.

**Results:** The result of the DPPH-free radical scavenging assay showed a synergistic antioxidant effects in CEOJ with an inhibition percentage of 71.39% higher than *J. tanjorensis* alone (66.80%) *O. gratissimum* alone (53.56%) and ascorbic acid (56%).

**Conclusion:** This results therefore inferred that the CEOJ produced better antioxidant activity than when used as a separate extract.

## Introduction

Nature has always catered for man as one of its own, be it in the provision of food, shelter and clothing, as flavours, as means of transportation and importantly in health. Plants have been used in traditional medicine for several thousands of years and their uses in different culture is well documented. (Johnson *et al*, 1999). It is the most common type of alternative medicine the world over (Ogboru *et al*, 2016). The reason for this is not far fetch, it is natural and as a result many regard it as also safe (Barnes *et al*, 2007). This trust has led to a paradigm shift from synthetic molecule to natural product inhibition in therapy. This was demonstrated by Newman *et al*, (2003) who showed that as at 2002, about 67% of new therapeutic agents were synthetic but as at the time of their report about five years later, only about 39% of new therapeutic agents were said to be synthetic while the rest are natural products. This huge shift alerted the World Health Organization (WHO) and 2008 they insisted that proper scientific investigations must be carried out in medicinal plants to ascertain their phytochemical constituent, their toxicity and the effectiveness or otherwise of the acclaimed medicinal uses. *Ocimum gratissimum* and *Jatropha tanjorensis* are among such plants that are increasingly used both for medicinal purposes and as food.

*O. gratissimum* belong to the family of Labiatae and it is the most abundant of the genus *Ocimum*. It is a perennial plant that is widely distributed in the tropics of Africa and Asia. It is commonly called Scent leaf, it is also called Tea bush or tree basil and in Nigeria locals call it different name depending on the tribe. For instance, it is called Nchuanwu in Igbo, Tanmota wangiwawang in Nupe, Daidoya-lagida in Hausa and Efinim in Yoruba. On the global scene it is called alfavata brava in Brazil, clavo in Cuba, Ajeka in India and Alfavacoa in Portuguese. Plant Resources of South-East of Asia (PROSEA) 2018 describes it as perennial shrub that grows to about 1-3m tall having an erect stem and vastly branched. Also leave opposite, petiole 2-4.5 cm long, slender, pubescent blade elliptical to ovate. The fruit consists of 4 dry 1-seeded nutlet enclosed in the persistent calyx and it is propagated by seed. It is usually cultivated for the production of its essential oils in the leaves and stems. Oils such as Eugenol, geraniol, linalool, citral and thymol have been extracted from it. These essential oil oils have reported important antimicrobial properties (Dubey *et al*, 2000).

The essential oils from *O. gratissimum* is being used in cosmetology and as a result, it has been vastly investigated for actions as antiinfectives. Pandey, (2017) showed its antibacterial and antifungal potential. It is also reported that the essential oil possesses etiological properties able to inhibit the virulent strains of *Shigella* isolates; the organism responsible diarrhea and reduces extracellular protese activity (Iwalokun *et al*, 2003). The hexane fraction of the extract also exhibited a very high antimicrobial activity against *Vibrio cholera* and *Klebsiella pneumonia* (Nweze & Eze, 2009). Pratheeba *et al*, (2015) reported the extract’s high efficacy against filariasis causing mosquito vector *Culex quinquifasciatus*.

*Jatropha tanjorensis* belong to the family of Euphoriceal and like *O. gratissimum*, it is a perennial shrub and it is also commonly distributed in Africa and Asia. It grows to about 1.8m tall and the leaves has granular hairs and are 3-5 lobed palmate. It is called Kattammanakku in India (Bharathy & Ulhayakumari, 2013). In Nigeria, it is called efo-iyana-ijapa by the Yorubas (Western Nigeria) and ugu oyibo in Igbo (Eastern Nigeria) and Chaya in Hausa (Northern Nigeria). It is commonly called hospital-too-far because of the general belief that it can cure many different diseases and anyone that takes it will rarely visit the hospital for treatment. Apart from its use as a vegetable used in food preparations, it has been reported to for its use in diabetes mellitus treatment because of its hypoglycemic property (Olayiwola *et al*, 2004). It has also been reported to possess blood cholesterol lowering ability (Oyewole & Akingbala, 2011) and thus useful in several cardiovascular diseases. Studies have demonstrated its antimicrobial effects (Iwalewa *et al*, 2005). The leaves of *J. tanjorensis* is widely consumed locally for treatment of anaemia and studies and study have shown that when the leaf powder was administered to rabbits, their hematological indices were greatly improved (Orheu *et al*, 2008).

The several health dangers posed by pro-oxidants which the human bodies encounter on daily basis and the frequent generation of reactive oxygen species (ROS) by the body systems has resulted in an increase in antioxidant research coupled with the fact that several diseases benefits immensely on treatment with antioxidant either as main treatment or as adjunct therapy, diseases such as cardiovascular diseases, cancer, immune system disorder and several degenerative diseases (Aruoma, 1998). Studies have also shown that antioxidants have successfully been used in management of aging process and age related illnesses (Sies, 1991). Several researches have been carried out on the antioxidant activities of *J. tanjorensis* as well as *O. gratissimum* for example Daniel *et al*, (2019) demonstrated the antioxidant activity of *J. tanjorensis* and its phenolic content and Sheneni *et al*, (2018) also reported the antioxidant activity of *O.gratissimum* but none has worked on combined antioxidant effects of these plants as both are mostly consumed together locally as food and in traditional medicine. This study therefore aims to investigate the combined antioxidant effect of *O. gratissimum* and *J. tanjorensis*.

## 2 Materials and Methods

### 2.1 Chemicals and Reagents

All the reagents and chemicals used in this research work were all of Analytical grade. The following reagents and chemicals were used in this study; DPPH (2,2-diphenyl-1-picryl hydrazyl), ascorbic acid (vitamin C), 95% ethanol, magnesium metal, distilled water, HCL (hydrochloric acid), ferric chloride, 2,4-diphenyl hydrazine, concentrated sulphuric acid, acetic acid, Meyer’s reagent ( potassium mercury iodide), 1,10-phenanthroline, ferrous sulphate (FeSO_4_), methanol, chloroform, sodium hydroxide, sodium nitroprusside, ethyl acetate, acetic acid and EDTA (ethylenediaminetetraacetic acid).

### 2.2 Preparation of Extracts

Fresh leaves of *J. tanjorensis* and *O. tanjorensis* were procured locally in Calabar and identification of the plants was done at the department of Botany of the University of Calabar. Separately, the leaves were thoroughly washed but carefully to avoid squeezing and kept to air-dry for about a week. Then the leaves were ground into powder using a blending machine. The ground samples were weighed and recorded for the calculation of percentage yield before soaking in methanol at room temperature for 48hours. At the end of the period, the mixture was filtered using Whatmann’s filter paper. This process was repeated two more times until the ground sample in methanol produced an almost colourless mixture. The residue was further soaked in methanol and kept for 24hours, after which the mixture was filtered and dried in mild temperature. The extract of *O. gratissimum* had a percentage yield of 6.1% while that of *J. tanjorensis* yielded 5.57%.

#### DPPH Free Radical Scavenging Assay

DPPH is the acronym for 2, 2-diphenyl-1-picryl hydrazyl, a well-known chemical in antioxidant research. It is a stable molecule and when dissolved in methanol it gives a deep violet colouration which is absorbed at the 517nm in spectrophotometer. The principle behind this technique is that antioxidants are able to react with the stable DPPH by giving electrons or hydrogen atom. This action reduces DPPH to DPPH-H or DPPH-R which is a substituted analogous hydrazine with a pale yellow colour and sometimes colourless solution as a result. This colour change can easily be monitored using a spectrophotometer (Emmanuel, 2021).

Since its inception by Blois in 1958, many scientists have remodeled and modified the DPPH model but for this particular study the method used was by Brand-William *et al*, 1995 and modified by Sanchez-Moreno 1998 with some modifications

## METHODOLOGY

Eight concentrations of the methanol extracts of *O. gratissimum* and *J. tanjorensis* as well as eight concentrations of the combined extracts in equal proportions were prepared, also 8 concentrations of ascorbic acid (standard) was prepared. The concentrations were done in order of serial dilution beginning from 250µg/ml to 1.9µg/ml (250µl/ml, 125µg/ml, 62.5µg/ml, 31.25µg/ml, 15.63µg/ml, 7.81µg/ml, 3.91µg/ml and 1.9µg/ml). Freshly prepared DPPH solution in methanol solution was kept in the dark awaiting usage. From the extracts preparations1ml each was measured out and put into a test tube, following this 1ml of the freshly prepared DPPH (152µM) was added to each of the test tube containing the extracts. The mixture was kept in the dark for 1 hour to set. The same procedure was repeated with concentrations of ascorbic acid and DPPH and also kept in the dark for 1 hour.

After 1 hour, their absorbance was read using a spectrophotometer at absorbance of 517nm. The process was also repeated for all the extracts and ascorbic acid. A blank solution of DPPH (without extract or ascorbic acid) and methanol was prepared and used to read the absorbance. The whole procedure was repeated three times to confirm the results. The free radical scavenging activity of the extracts and ascorbic acid was calculated from the absorbance values as percentage inhibition using the equation:

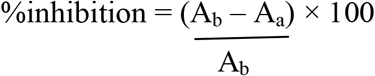

Where A_b_ = Absorbance of blank

A_a_ = Absorbance of extract or ascorbic acid.

## Results

DPPH Free Radical Scavenging Activity.

The methanol extracts of *O. gratissimum* and *J. tanjorensis were* tested for their free radical scavenging ability against DPPH, a stable compound and their percentage inhibition was calculated as reported below.

### 3.2.1 Percentage inhibition of *O. gratissimum*

The percentage inhibition of *O. gratissimum* was observed to be between 37.9% and 61%. The highest inhibition percentage was observed at the concentration of 250µg/ml and it was significantly different from the inhibition percentage of the lowest concentration (1.9µg/ml) which stood at 37.9%. Following the lowest concentration was the 3.91µg/ml concentration with an inhibition percentage of 41.90%. There was no significant difference in percentage inhibition of the concentrations of 7.81µg/ml, 15.63µg/ml and 31.25µg/ml with regards to their inhibition percentage at 45.05%, 46.26% and 50.20% respectively. Furthermore, there was no significant difference in the percentage inhibition at the concentration of 62.50µg/ml and 125µg/ml with their percentage at 56.91% and 58.89% respectively. Another important factor worthy of note is that there was dose-dependent increase in the inhibition percentage values of the extract of *O. gratissimum* as well as the control (ascorbic acid).

### 3.2.2 The percentage inhibition of *J. tanjorensis* Extract

The percentage inhibition of the methanol extract of *J. tanjorensis* showed the highest concentration of the extract (250µg/ml) recorded the highest inhibition percentage (90.12) and this was significantly different from the rest of the concentration on statistical analysis. The lowest concentration of the extract (1.9µg/ml) gave the lowest inhibition percentage but there is no significant difference between the inhibitions of this lowest concentration and the next concentration (3.91µg/ml). The extract concentrations at 1.91µg/ml, 3.91µg/ml, 7.8 µg/ml, and 15.63µg/ml have no significant difference in the values of their percentage inhibition upon analysis. The inhibition percentage of the above concentrations though not significantly different, the increase was concentration-dependent.

### 3.2.3 The Percentage Inhibition of Combined Extracts of *O. gratissimum* and *J. tanjorensis*

One of the major aims of this research was to determine the antioxidant activity of the combined extracts of *O. gratissimum* and *J. tanjorensis* as both leaves are commonly consumed together as vegetables. The combined extracts showed a higher antioxidant activity when compared to the activity of the individual plants. For instance, the lowest concentration (1.91µg/ml) of the combined extract showed a percentage inhibition value of 67.81%, this is significantly different from the inhibition percentage of the lowest concentration of *O.gratissimum* (1.91µg/ml) with a value of 37.99% which is also significantly different from the inhibition percentage value at the lowest concentration of *J. tanjorensis* (1.91µg/ml) with the value of 59.68%. There was a significant difference in the percentage inhibition value at the concentration of 7.81µg/ml with the rest of the concentrations. Again the rate of increase in the percentage inhibition values was not in correspondent with the increase in the concentration of the extract meaning that it is not concentration/dose dependent.

**Figure 1.**
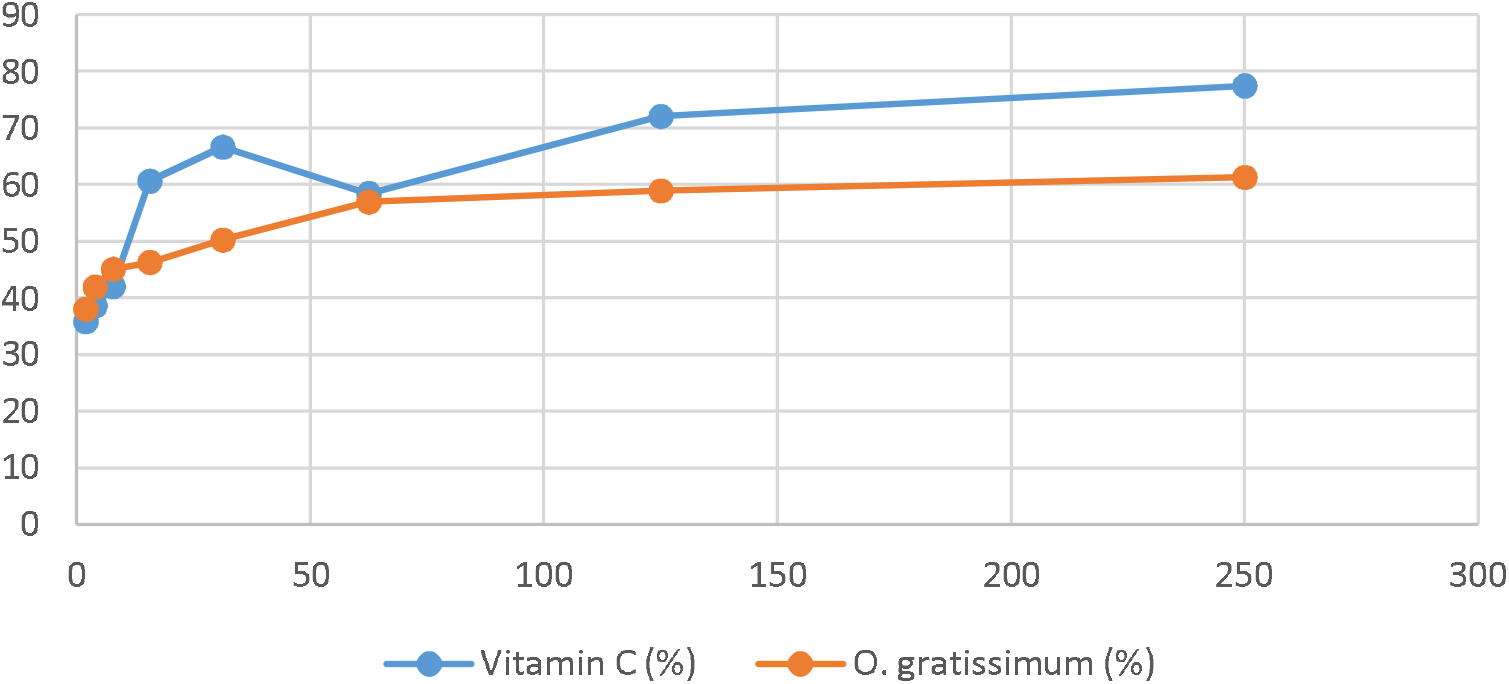
Percentage Inhibition of O.gratissimum and Vitamin C.

**Figure 2.**
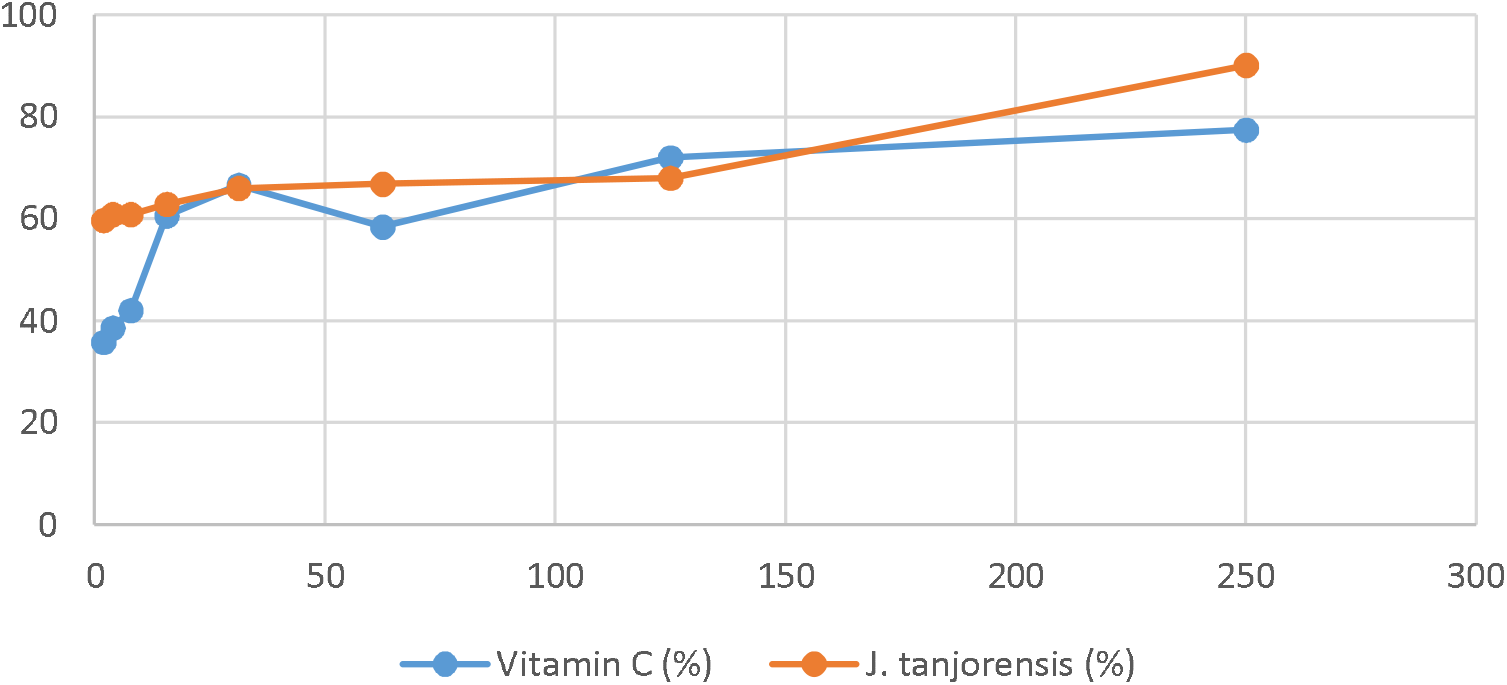
percentage inhibition of vitamin C and J. tanjorensis

**Figure 3.**
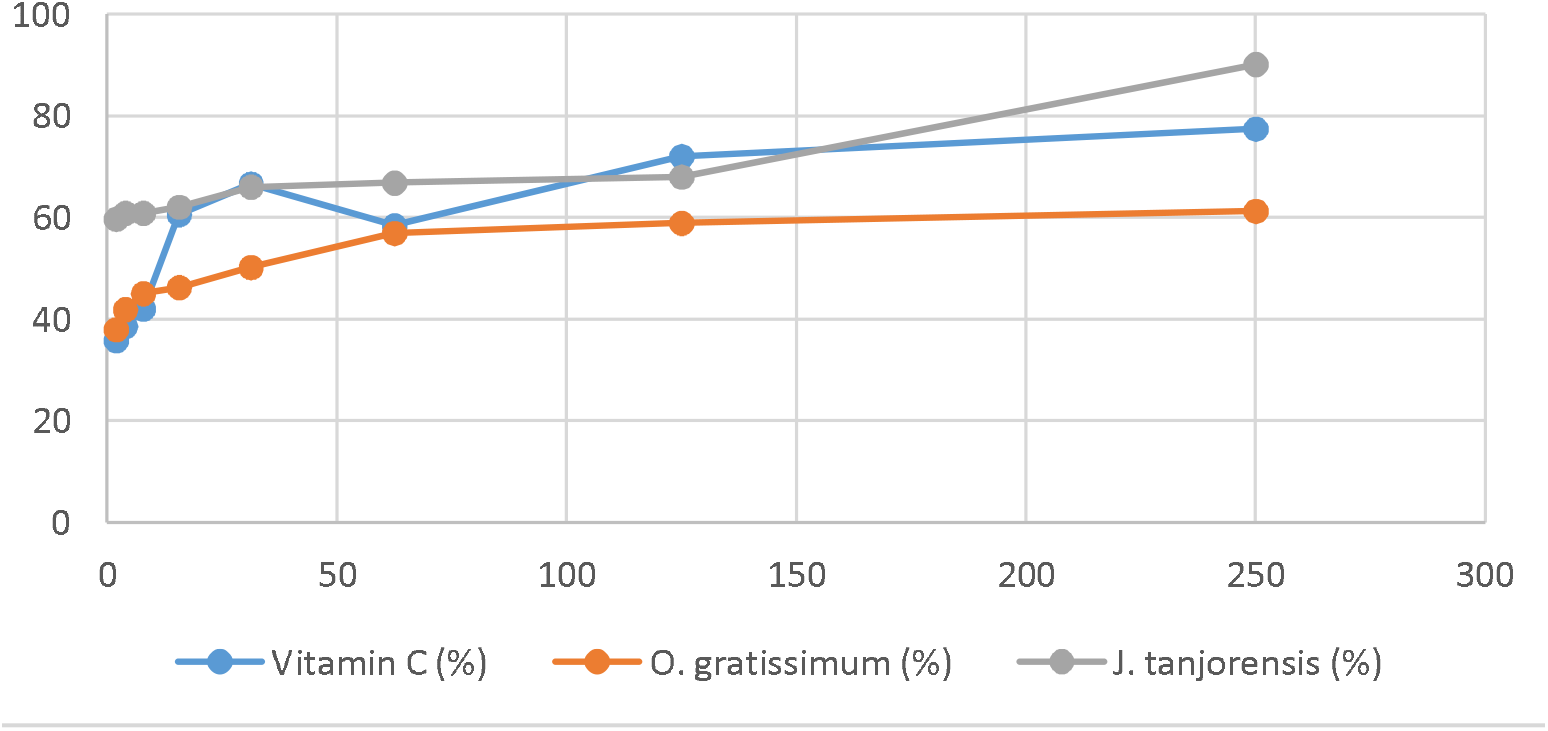
Percentage Inhibition of Vitamin C, O. gratissimum and J. tanjorensis

**Figure 4.**
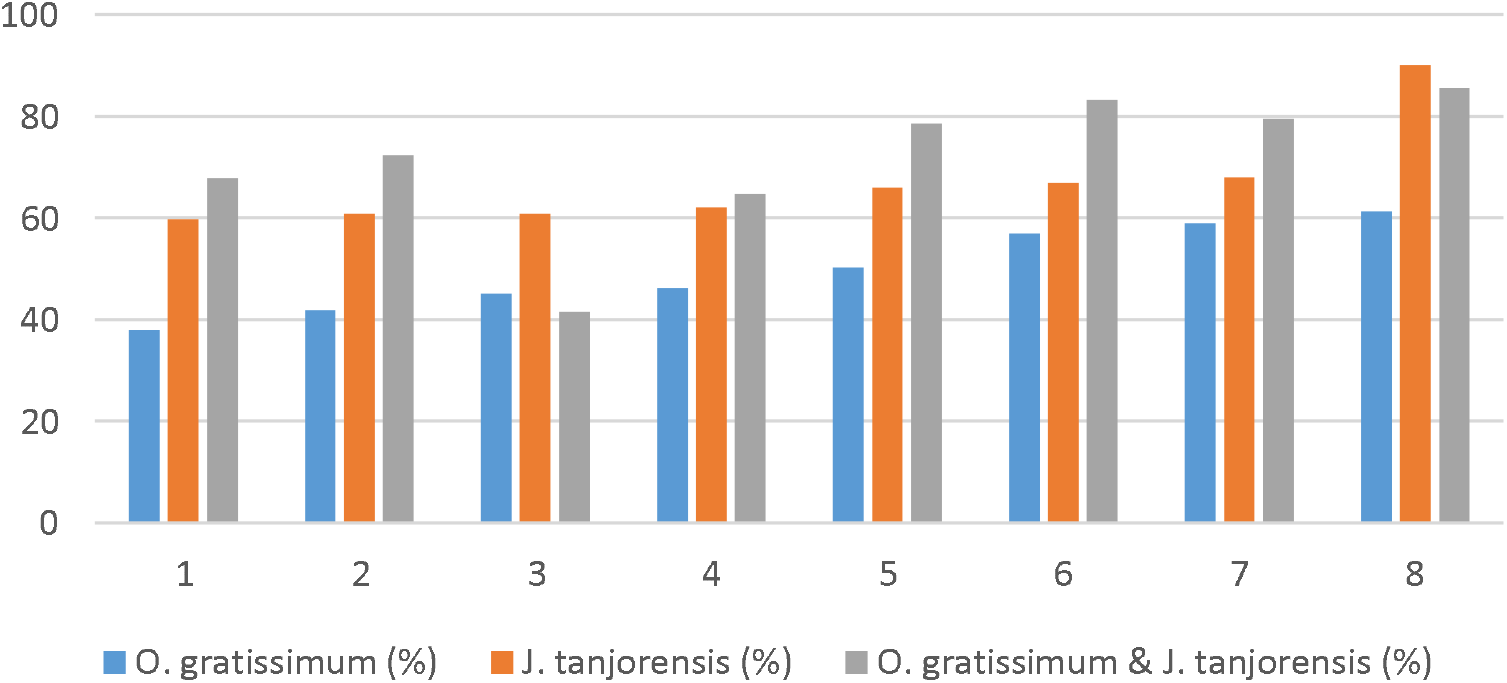
Percentage inhibition of O. gratissimum, J. tanjorensis and both extracts combined.

## Discussion

In the investigation for antioxidant activity of the extracts, the DPPH free radical scavenging method was used for the research. The method is based on measuring the scavenging capacity of antioxidant agents against the DPPH; whose odd electron of nitrogen atom is reduced by receiving a hydrogen atom from the antioxidant to the corresponding hydrazine (Contreras-Guzman & Strong 1982). The DPPH free radical scavenging assay was originated by Bois in 1958, it offers the first approach for evaluating the antioxidant potential of a compound (Kedare & Singh, 2011), and since then it has been modified by different scientists to meet different scientific needs.

A major advantage of the DPPH assay compared to other antioxidant models is the fact that it is highly and easily reproducible, it is very simplified and it also measures the overall antioxidant activity Parkash, (2011). It can also measure the antioxidants of fruits and vegetables (Sendra *et al*, 2006). Prior *et al*, (2005) reported that it can be used to measure both lipophilic and hydrophilic substances. Another important advantage according to Bondet *et al* (1997), is that the effectiveness of the antioxidant is measured in ambient temperature, this eliminates the risk of thermal degradation of the tested substance and the result gotten from the assay is comparable to other free radical scavenging assays (Gil *et al*, 2000).

However, despite its numerous advantages the DPPH free radical scavenging method of testing antioxidants has its own limitations. The first is that the absorbance of DPPH in methanol and acetone decreases under light (Min, 1998), this makes it difficult to get the accurate absorbance of the substance as most of the absorbance is read under light. Secondly, there is constant interaction between DPPH radicals and other radicals present. Even the time-response curve to reach the steady state is not linear with different ratios of antioxidant/DPPH (Brand-William *et al*, 1995). Sanchez-Moreno *et al* (1998) saw this as a very big disadvantage insisting that if the results are to be graphically represented or presented in bar charts even if the same data is available in nominal form, it will pose a great difficulty.

For this present study, the DPPH free radical scavenging method used was by Brand-William *et al*, (1995) and modified by Sanchez-Moreno *et al*, (1998) with some modifications. The blank (DPPH and methanol) at absorbance of 517nm gave a value of 0.253 which is less than 1.0 to conform to what Bois (1958) reported that for an experiment to be valid, the initial DPPH concentration should produce an absorbance value of less than 1.0. From the various absorbance value of the extract and standard, their percentage inhibition of the DPPH free radical was calculated using the percentage inhibition method. The percentage inhibition of the extract of *O. gratissimum* ranged from 37.90% to 61.26%, that of *J. tanjorensis* from 59% to 90% while the combined extracts ranged from 41% to 85%. According to a study by Molyneux (2004), an agent is said to possess potent antioxidant activity if the percentage inhibition of the said substance is above 50%. This according to the study results from the fact that antioxidants foods and plant materials like extract maybe water soluble, fat soluble or insoluble or bound to cell walls and so may not be readily free to react with DPPH all at the same time. Therefore, if in spite of all the above mentioned, the agent is still able to produce an inhibition percentage of 50% and above, such an agent is a potent antioxidant agent the study concluded.

## Conclusion

Overall, the combined extracts of *O. gratissimum* and *J. tanjorensis* produced the highest inhibition percentage with a mean inhibition of 71.76%, this was followed by the extract of *J. tanjorensis* with a mean inhibition of 66.90% while the methanol extract of *O. gratissimum* yielded a mean inhibition of 49.81%. The control (vitamin C), yielded a mean inhibition of 56.1%. This may justify the rationale behind the combined consumption of theses extracts as they are more effective together than when used individually in terms their DPPH free radical scavenging ability.

